# Optimizing sequential gene expression modulation for cellular reprogramming - Coupled Boolean modeling and Reinforcement Learning based method

**DOI:** 10.1101/2024.03.19.585672

**Authors:** Vivek Singh

## Abstract

Regenerative medicine entails regenerating damaged cells and tissues from healthy cells through the process of cellular reprogramming. The most common method to achieve reprogramming of cells of one type to another is by modulating the activity of specific genes. However, identifying the most suitable genes for reprogramming is a challenge due to their large number in humans and their complex interactions.

This study describes a computational method to predict sequential gene expression modulation for reprogramming a starting cell type to a target cell type. The proposed method integrates: (1) a Boolean model of the concerned gene regulatory network (GRN); and (2) a reinforcement learning (RL) based model for optimization. The Boolean model is used to capture the dynamic behavior of the GRN and to understand how the gene expression modulation alters its behavior. RL model is used to optimize sequential decision-making of predicting the suitable sequence of gene expression modulation.

Coupling of the Boolean model and RL plays a crucial role in the proposed computational method. Boolean model captures the GRN dynamics, and thereby, constrains the combinatorially large state space. The RL model operates in this constrained state space and uses the Boolean model to evaluate the effect of modulations on GRN dynamics to predict the sequence of suitable gene expression modulations.

Applicability of the proposed method is demonstrated using a toy network of 4 genes, and a biological network of heart development representing the dynamics of 15 genes.

## 1. Introduction

Regenerative medicine is an evolving branch of medicine that deals with repairing or replacing damaged and diseased cells, tissues and organs with the healthy ones [1], [2]. The first step towards this is the regeneration of healthy cells of specific types that can develop into required tissues and organs. This is achieved through the process of cellular reprogramming [3], by: (1) cellular differentiation of pluripotent cells [4], [5], [6] or (2) transdifferentiation of somatic cells directly to the target cell type [7]. The latter has been shown to be more efficient and has reduced risk of teratoma formation [8].

Cells maintain their identity, subtype and phenotypic properties through characteristic gene expression profiles, regulated by the dynamic activities of transcription factors (TFs) and their corresponding target genes [9]. In accordance with this, targeted cellular reprogramming has been achieved by modulating the expression of specific genes (generally TFs) [8], [10]. TFs regulate the expression of other (target) genes through complex gene regulatory networks (GRNs), and establish phenotype-determining gene expression profiles in the cells [11]. For targeted cellular reprogramming, the gene expression modulations can be done simultaneously at the initiation, or more recently in a sequential manner in concordance with the gene expression profiles of the starting and target cell types. Sequential gene expression modulation has proven to be more efficient in reprogramming and/or in establishing the maturation of the target cell type [12], [13].

Although cell-type determining TFs are known to some extent, identifying the correct TF(s) that can serve as a switch to induce targeted cellular reprogramming is a challenge. This is, in part, due to the large number of protein-coding genes and TFs in them (∼19,000 [14] and ∼16,00 [15] respectively in humans), and the complex gene regulatory networks that regulate their functional interdependence. The large number makes identification of suitable TFs difficult, and the complex functional interdependence complicates the matter further because gene expression modulations can lead to cascading effects on the gene expression profile making it difficult to assess the effect of any such modulation. Therefore, gene expression modulation for cellular reprogramming is still largely empirical or based on trial-and-error, resulting in low efficiency of reprogramming [6], and tumorigenicity [5] and immune rejection [2], [5] of the target cells in the body. To overcome these challenges, computational methods that can exploit available biological data and data-driven approaches are needed. Such methods can predict suitable gene expression modulations *a priori*, and supplement and guide the choice of genes for specific cell type reprogramming [16].

The prediction of gene expression modulation for targeted cellular reprogramming is naturally a sequential decision-making problem wherein genes to be modulated and their sequence need to be predicted. Optimizing such sequential decision-making has recently seen a lot of interest across many disciplines [17]. Although, it has long been studied in control theory and computer science [18], the advent of deep learning and recent successes of reinforcement learning in solving complex problems [19], [20], [21] has led to an explosion of interest on this topic.

Gene regulatory networks and their dynamic behavior play an important role in defining cellular gene expression profile, and thereby, their identity and phenotype [9]. Thus, it is important to account for them while developing a computational method for predicting suitable gene expression modulation. Systems Biology offers multiple formalisms to gain holistic understanding of such complex regulatory processes [22], [23], [24]. Boolean network modeling is one such formalism that approximates the regulatory interactions through binary activity levels (0: repression and 1: activation), allowing it to model large-scale biological networks such as gene regulatory networks [22].

A few existing computational methods have leveraged dynamic behavior (or at least network topological properties) of GRNs with or without control theory for predicting gene expression modulation for targeted cellular reprogramming. For instance, NETISCE [25] used a combination of signal flow analysis [26], feedback vertex set control [27] and machine learning to predict the required gene expression modulation; Mandon *et al.* [28] used static analysis of the Boolean network’s interaction graph for predicting reprogramming strategies; and Cifuentes-Fontanals *et al.* [29] used model checking on Boolean network model of GRN for predicting the reprogramming strategies.

However, to the best of authors’ knowledge, none of the existing methods combine the strength of reinforcement learning for optimizing sequential decision-making along with a model capturing the dynamic behavior of the concerned GRN. This integrated method holds potential for predicting sequential gene expression modulations that are biologically feasible and bounded by relevant constraints of the concerned GRN and its dynamics.

This study proposes a computational method that utilizes the dynamics of GRNs predicted using Boolean network model and couples it with a Q-learning based reinforcement learning model for predicting optimal sequential gene expression modulations for targeted cellular reprogramming. The capability of the method is demonstrated through a toy network as well as a biological network corresponding to early heart development (referred to as cardiogenesis model henceforth).

## 2. Background

### 2.1 Boolean network modeling

#### 2.1.1 Boolean network models

Boolean network models are a special case of discrete dynamic models, and have been commonly used for modeling dynamical behavior of gene regulatory networks [24], [30], [31]. In a Boolean network, nodes represent the components (e.g. genes) of the modeled system and the edges represent regulatory interactions between the components. Each node has one of the two possible Boolean states - 0 (repressed) and 1 (activated) representing the biological state of the designated component [32], [33]. These binary states are a rough approximation of the states of biological components [34]. For instance, for a gene as a node, the two states can correspond to the presence or absence of that gene product.

States of the nodes are determined by Boolean state transition functions that are dependent on the states of other nodes and the regulatory interactions between them [24], [30]. These state transition functions map the state of the Boolean network to a succeeding state, and allow a Boolean network model to exhibit dynamical behavior in simulations [34].

State of a Boolean network is defined by the states of each of its nodes, and is represented by a binary vector of length n (n = number of nodes in the network). Elements in this binary state vector represent the states of nodes (i.e. 0 or 1 corresponding to their repressed or activated states). As the state of each node is represented by one of the two possible values, there is a finite number of possible states (called as state space) for a Boolean network. For n nodes, this is equal to 2^n^ possible states.

The dynamics of a Boolean network model is simulated by incremental execution of one or more of its state transition functions. They change the states of corresponding nodes in a series of discrete time steps called transitions [31], [34]. The spectrum of all possible transitions is illustrated using the state transition graph, wherein the nodes represent the states of the Boolean network and the edges represent the transition between states [35].

The order in which the state transition functions are executed during the dynamic simulation is governed by the updating scheme [55]. Most frequently used updating schemes are synchronous and asynchronous. Synchronous scheme updates the state of all nodes of the Boolean network at the same time and is deterministic in nature. Contrarily, an asynchronous scheme updates randomly the state of a single node per transition, making each execution non-deterministic [35], [36].

#### 2.1.2 Attractors in Boolean network models

Simulation of Boolean network models can lead to stable dynamic behavior characterized by periodic sequences of states, called attractor states (or simply attractors) [31], [34]. Once reached, the Boolean network stays in the attractor state unless an external perturbation occurs [35]. These attractors represent long-term behavior of Boolean networks and are interpreted as biological phenotypes or physiological endpoints (e.g. specific types of cells, or physiologically distinct states of cells of the same type) [37], [38]. Attractors can be subdivided into different types. Steady state attractors comprise only one Boolean network state. Additionally, there are attractors with more than one state, for example, simple cycles, which are formed by a sequence of states of a certain length which are periodically repeated [24], [39]. For more details and other types of attractors, refer: [39], [40], [41].

#### 2.1.3 Applications

In terms of complexity, Boolean network models lie between static network models and continuous dynamic models, making them a tractable and powerful approach to model large-scale biological systems. Boolean network models can be used to describe the qualitative temporal behavior of the system and to understand how perturbations may alter its behaviors [31]. Boolean network models have been successfully applied in modeling many gene regulatory and signaling networks in a variety of organisms [42], [43], [44], [45], and have been reviewed in several articles [46], [47], [48].

### 2.2 Reinforcement learning and Q-learning algorithm

#### 2.2.1 Reinforcement learning

Reinforcement Learning (RL) is a machine learning technique to learn sequential decision-making in complex scenarios [49]. RL is inspired by trial-and-error based human/animal learning, wherein the learning is driven by a scaler quantity (the ‘reinforcement’ or the ‘reward’) and the goal of the algorithm is to maximize the expected future cumulative reward [50].

RL is formulated as an optimal sequential decision-making algorithm with the ability to account for stochasticity within the system [51]. In RL, the learner is referred to as the agent and the world in which the agent lives and interacts is referred to as the environment. The environment can be in any one of the states defined for a particular model. Similarly, an agent can take a defined set of actions in a particular model. Moreover, a formal description of an RL problem also includes a transition function that describes how the environment will respond (i.e. change its state) to the agent’s actions, and a reward function that defines how good (or bad) the action was [50].

The goal of the RL agent is to take actions that maximize the expected future cumulative reward (i.e. the sum of the rewards during an episode or the entire life of the RL agent depending on the task). The actions taken by the agent are guided by either trial-and-error, or based on the state of the environment and rewards, or a combination of both. By performing these actions iteratively, the agent learns an optimal behavioral strategy based on the rewards received from previous interactions [49].

The decision-making process in RL is formalized as a Markov Decision Process (MDP) [51]. MDP is a formulation of sequential decision-making where both immediate and future rewards are considered [18]. MDP provides a formalism to the decision maker (agent in our case) for rationalizing the planning and action in the face of uncertainty, i.e. it evaluates the best action the agent should take considering the state of the environment [51].

#### 2.2.2 Q-learning algorithm

Q-learning is a model-free, off-policy reinforcement learning algorithm, which is used to find the optimal action-selection policy i.e. the best course of action in a given state [17]. The ‘Q’ in Q-learning usually stands for quality, as the algorithm calculates the maximum expected rewards for a given action in a given state [52]. It does not need a model of the environment (hence the term “model-free”), and it can deal with stochastic transitions and rewards without the need for adaptations.

Q-learning uses an off-policy control i.e. the behavior policy and the target policy are different [51]. For any finite Markov decision process, Q-learning can find an optimal (action-selection) policy given infinite exploration time and a partly random policy using Bellman optimal equations and the 𝞊-greedy policy [51], [53]. Unlike other RL algorithms, Q-learning has simple Q-functions, hence it has become the foundation of many other RL algorithms [53].

#### 2.2.3 Applications

RL has emerged as an efficient technique for solving complicated sequential decision-making tasks in recent years. It offers a great opportunity to open new frontiers in technology where system models are unavailable or too difficult and/or expensive to build [49]. Its applications range from robotics [54], [55], [56] to games such as Go, chess [57], [58], and even complex video games such as StarCraft II [59].

## 3. Materials and methods

The proposed method couples a Boolean network model with a Q-learning based RL model to predict optimal gene expression modulation for targeted cellular reprogramming. Two such coupled models are developed, one using Boolean network model of a toy network and the corresponding RL model, and the other using Boolean network model of cardiogenesis core gene regulatory network and the corresponding RL model. A schematic diagram of the proposed method is shown in Figure 1.

**Figure 1:**
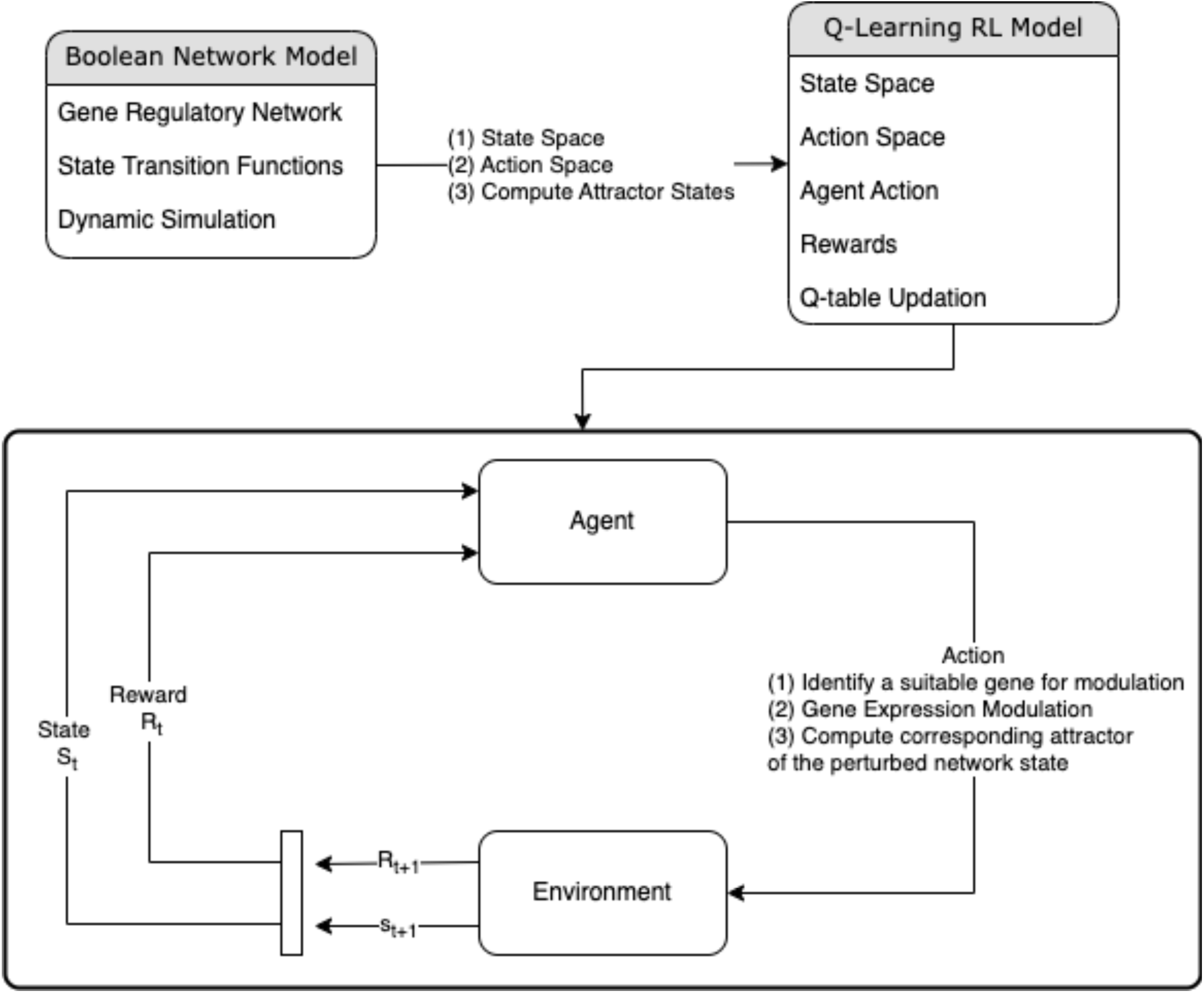
Schematic representation of the proposed method. Top boxes indicate the roles played by the Boolean network model and the Q-learning based RL model in the proposed method: Boolean network model defines the state and action spaces for the RL model and computes the attractor states corresponding to the perturbed network state; Q-learning based RL model defines the agent actions, rewards and Q-table updation strategy. Bottom box depicts the interaction between agent and environment, and the resulting state alterations and rewards. The steps involved in the agent’s action are also listed.

### 3.1 Boolean network models and their dynamics

A toy model and a murine early heart development (cardiogenesis) model [60] are used in this study. They are briefly described below, and the detailed information on nodes, interactions and the Boolean state transition functions are available in Supplementary data.

The toy model consists of 4 hypothetical nodes and the interactions between them. The cardiogenesis model represents the dynamics of the known core gene regulatory network involved in early heart development in mice. The model is based on published temporal gene expression data of relevant transcription factors and growth factors and includes known regulatory interactions [60]. As detailed in section 2.1.1, the state of each node in the network can take binary values: 0 and 1 representing repressed or activated states respectively of the corresponding gene. The state space of the Boolean network model, for instance, the cardiogenesis model, has 2^n^ = 32,768 possible states, with each state represented by a binary vector of length n = 15 (n = number of nodes in the network).

The dynamics of the Boolean network models are simulated using asynchronous update scheme as it is considered to be a closer representation of the biological scenario for gene regulatory network dynamics [61]. The simulations serve to capture the dynamical behavior of the concerned gene regulatory networks, and identify their attractor states (see section 2.1.2 for details).

Attractor states (or attractors) represent stable dynamic behavior of Boolean network (see section 2.1.2) and have been linked to biological phenotypes [31], [34], [37], [38]. For example, attractors can characterize distinct cell types, or physiological states of the same cell types. Thus, cellular reprogramming of a cell type or state to another can be viewed as transitioning from one attractor state (starting cell type or state) to another attractor state (the desired target cell type or state).

All the computations and simulations related to these Boolean network models are performed using PyBoolNet [62].

### 3.2 Q-learning based Reinforcement Learning models

In this study, Q-learning algorithm is used to develop the RL models for predicting sequential gene expression modulations for reprogramming an attractor in the respective Boolean network model to the desired attractor state. The RL models utilize the Boolean network models to: (1) determine their state and action spaces; and (2) predict how the gene expression modulation performed by the RL models affect the attractor states of the respective networks.

#### 3.2.1 RL environment

Environment of a RL model is the world context in which the RL agent operates. The agent interacts with the environment by taking actions, receiving rewards and transitioning to new states based on its actions. The environment provides the agent with its initial state, the current state and the updated states after each action. It also determines the reward associated with each state-action pair. A custom RL environment is created using Gymnasium [63] to address the needs of this study.

##### 3.2.1.1 State and action spaces

The RL environment is built with the state space and action space defined by the corresponding Boolean network model. Essentially, this means that, for instance, for cardiogenesis model, the RL environment can be in any of the 2^n^ = 32,768 possible states of the Boolean network, and the RL agent can perform expression modulation of any of the n = 15 nodes (genes) in the Boolean network (action space). Moreover, the initial and the desired target state of the environment are set during model definition.

##### 3.2.1.2 RL agent actions and the resultant alteration in RL environment states

In this study, an action by RL agent involves the following: (1) selecting a gene from the set of all the genes in the network; and (2) flipping the state value of the selected gene i.e. if a gene is in a repressed state (= 0), it would be flipped to activated state (= 1), and vice versa. In biological terms, this action represents predicting a suitable gene and modulating its expression to either activation or repression.

The action by RL agent results in a perturbed state of the environment i.e. of the Boolean network wherein the element corresponding to the modulated gene has its value flipped. Boolean network model is then simulated with this perturbed state and transitioned to the corresponding attractor state. This attractor state of the Boolean network becomes the new state of the RL environment as a result of the action by the RL agent.

##### 3.2.1.3 Rewards

The RL agent (i.e. the model) learns on the basis of the rewards it receives from the environment based on its actions and the corresponding alterations in the state of the environment. In this study, the reward is defined as a function of the Hamming distance [64] between the current state and the target state of the environment (see Equation 1). This Hamming distance is used to assess how far is the current state of the environment from the target state. A distance of zero indicates that the current state is the same as the target state i.e. the goal is achieved.

An action leading to smaller Hamming distance (i.e. bringing the current state closer to the target state) receives higher reward and vice versa. The model provides dense rewards i.e. the agent receives reward at each time step.

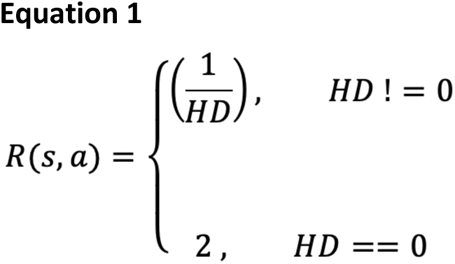

where R(s,a) = Reward for taking action **a** at state **s**, HD = Hamming Distance between the current state and the target state of the environment

#### 3.2.2 Q-learning algorithm

As detailed in section 2.2.2, Q-learning is a model-free and off-policy RL algorithm that learns suitable action to be taken by the agent in a given state that maximizes the expected cumulative reward (i.e. the sum of rewards during an episode or the entire life of the RL agent, depending on the task) [52], [65]. The Q-learning algorithm utilizes a Q-table, representing the state-action values. For each state, the Q-table stores the maximum expected future reward of each action at that state. The Q-values in the Q-table are used to determine the best actions for each state in the RL environment [65]. The steps involved in the Q-learning algorithm are discussed in the below sub-sections.

##### 3.2.2.1 Initializations

Initial state of the RL environment state is chosen randomly from the state space. This state may or may not be an attractor state. The Q-table is initialized with all zeros implying that, at any state, all of the possible actions are equally favorable.

##### 3.2.2.2 Selecting and performing an action

The RL agent chooses the most favorable action **a** for the current state **s** based on the current Q-values. Since all the Q-values are zero at the start, the action selection is random. As the agent continues to interact with the environment, the values in the Q-table get updated (detailed in section 3.2.2.3) and the action selection is guided by the updated Q-table. Equation 2 represents this action selection approach, and is used to guide the action selection at each step.

This action selection approach is based on the concept of exploration / exploitation [18]. Exploration indicates selecting actions randomly allowing exploration of the environment, and is relevant at the beginning of model training when the agent does not know anything about the environment (i.e. the Q-table has all zero values). On the other hand, exploitation refers to selecting the best action based on the current values in the Q-table i.e. exploiting the current information stored in the Q-table. Thus, the action selection relies on exploration at the beginning, which progressively decreases as the Q-table gets updated with the experience of the agent interacting with the environment.

The steps involved during an action are detailed in section 3.2.1.2. Each action results in altering the state of the environment and the agent receives a reward for its action as detailed in section 3.2.1.3. Based on these, the Q-table is updated as discussed in section 3.2.2.3.

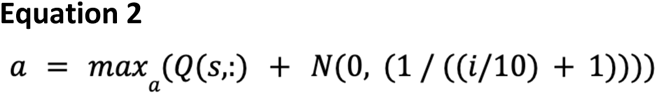

where, a = Selected best action; Q(s,:) = Q-values for all the actions corresponding to state, s; i = Episode number

##### 3.2.2.3 Q-table updation

After the completion of each step i.e. an action by the RL agent, the resultant alteration in the state of the environment and the reward received by the agent, the Q-table is updated. The respective Q-value in the table is updated based on the current value, the reward received by the agent for the action it took in a particular state, and the values of the hyperparameters defined in the model (see Equation 3). Equation 3 is formulated on the basis of the Bellman optimality equation [18].

There are two main hyperparameters in the model, the learning rate 𝛼, and the discount factor 𝛾. Learning rate determines the extent to which the newly acquired information by the agent during the current step overrides the old information. A low learning rate indicates that old information is preferred over new information and vice versa. Discount factor determines how much the future rewards are valued [9]. A low discount factor indicates that the algorithm prefers rewards in the short term, while a high discount factor indicates that the future rewards are valued more [65]. For this study, 𝛼 = 0.1, and 𝛾 = 0.95.

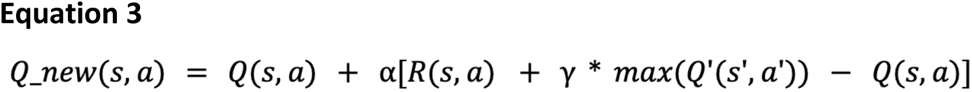

where Q_new(s,a) = New Q-value for state **s** and the action **a**; Q(s,a) = Current Q-value for state **s** and the action **a**; ɑ = Learning rate; R(s,a) = Reward for taking action **a** at state **s**; γ = Discount rate; max(Q’(s’,a’)) = Maximum expected future reward given the new state **s’** and all possible actions at that new state

### 3.3 Model training and inferencing

The coupled Boolean and RL models corresponding to the toy network and the cardiogenesis network are trained in a similar way, as described below.

The steps described in section 3.2.2 are performed iteratively for training the coupled models. As is detailed in section 3.2, the Q-learning models are coupled with the corresponding Boolean network models. The resultant updated Q-tables after model training are then used for inference from the models.

Both the coupled models are trained for 300 episodes. Each episode begins with a random state of the RL environment (i.e. the state of the corresponding Boolean network). The episode continues till the goal is achieved i.e. environment reaches the target state (Hamming distance between current state and the target state = 0), or maximum number of steps (=100 in this study) are exhausted. At each step, if the environment is not at the target state, the agent takes an action based on Equation 2, resulting in an updated state of the environment and the associated reward to the agent.

For performing inference from the trained model, the updated Q-table is used. The goal being to find the optimal sequential gene expression modulation to reach the target attractor state from any state in the state space of the RL environment (i.e. the state space of the corresponding Boolean network). The inferencing episode continues till the agent reaches the target state or exceeds the threshold of steps (=100 in this study).

## 4. Results

### 4.1 Construction of the models

The proposed method couples a Boolean network model with a Q-learning based RL model to predict optimal gene expression modulation for targeted cellular reprogramming. In this study, cellular reprogramming of a cell type or state to another is represented as transition from one attractor state (starting cell type or state) to another attractor state (the desired target cell type or state) (see section 3 for details). Two coupled models are developed, one corresponding to a toy network and the other of cardiogenesis core gene regulatory network. The results corresponding to the cardiogenesis model are discussed in the main text; results of the toy model are available in Supplementary data.

#### 4.1.1 Boolean network model contributes the biological context and constraints in the proposed method

In the proposed method, Boolean network model contributes to the following aspects: (1) representing the interactions among genes and the dynamic behavior of the concerned GRNs; (2) identifying biologically relevant states of the GRN networks (i.e. attractor states); and (3) predicting the effect of gene expression modulation on GRN states. Most importantly, given that the attractor states are linked to biological phenotypes [34] such as cell types or physiological states of a cell type, Boolean network model helps map the dynamics of GRNs to the objectives of cellular reprogramming.

#### 4.1.2 Q-learning based RL model predicts the sequential gene expression modulations for targeted cellular reprogramming

The corresponding Q-learning based RL model is used to solve the sequential decision-making problem i.e. predicting the genes, their type of modulation (i.e. activation or repression) and the sequence of modulation that would reprogram (i.e. transition) the respective Boolean networks from a starting state to a desired target attractor state. The RL model is coupled with the Boolean network model, which predicts how the gene expression modulations performed by the RL model affect the states of the network, thereby providing biological context in this coupled model.

The resultant trained coupled model is used for inference i.e. to predict the required gene expression modulations for targeted cellular reprogramming, represented here as transition from one attractor state to a target attractor state.

### 4.2 Cardiogenesis coupled model and its application in predicting the gene expression modulations for transitioning to FHF state

#### 4.2.1 Model development and training

##### 4.2.1.1 Boolean network model of cardiogenesis core gene regulatory network

The model contains 15 nodes (genes) and the interactions among them represented as Boolean state transition functions for each node (refer Supplementary data for details). Key signals from other tissues that are required for heart development are represented by the following four nodes (genes): exogen_BMP2_I, exogen_BMP2_II, exogen_CanWnt_I, exogen_CanWnt_II. exogen_BMP2_I and exogen_BMP2_II represent inputs for non-cardiac BMP2 signaling, and exogen_CanWnt_I and exogen_CanWnt_II represent inputs for non-cardiac canonical Wnt signaling [60]. Moreover, the node, exogen_BMP2_I was kept always active in the original publication [60], but is allowed to change value and behave as a self-loop as done in [28].

Given 15 nodes in the network, the state space has 2^15^ = 32,768 possible states. Of these, 6 states are predicted to be the attractor states by the Boolean network mode. Two of them are potential artifacts of the model, one with all the genes repressed and other with only BMP2 and its exogene activated. The attractors, First Heart Field (FHF) and Second Heart Field (SHF), represent the key phenotypes of heart development [60]. The remaining two are close to either FHF or SHF, with only a few genes having different values. These are termed ‘FHF muted’ and ‘SHF muted’ respectively with both having exogen_BMP2_I repressed. The state vectors for attractor states indicating the active and repressed genes are shown in Table 1.

**Table 1:** Attractors in the cardiogenesis model and the corresponding state vectors of the Boolean network. Rows: Attractor states; Columns: Genes in the network; 0: Gene is repressed; 1: Gene is active

|  | Bm<br>p2 | Dkk1 | Fgf8 | Foxc1<br>_2 | GATA<br>s | Isl1 | Mesp<br>1 | Nkx2_<br>5 | Tbx1 | Tbx5 | CanW<br>nt | exoge<br>n_BM<br>P_I | exoge<br>n_BM<br>P2_II | exoge<br>n_Can<br>Wnt_I | exoen<br>_can<br>Wnt_I<br>I |
| --- | --- | --- | --- | --- | --- | --- | --- | --- | --- | --- | --- | --- | --- | --- | --- |
| Inactive | 0 | 0 | 0 | 0 | 0 | 0 | 0 | 0 | 0 | 0 | 0 | 0 | 0 | 0 | 0 |
| Active<br>BMP | 1 | 0 | 0 | 0 | 0 | 0 | 0 | 0 | 0 | 0 | 0 | 1 | 1 | 0 | 0 |
| SHF<br>muted | 0 | 1 | 0 | 1 | 1 | 1 | 1 | 1 | 1 | 0 | 1 | 0 | 0 | 1 | 1 |
| SHF<br>state | 0 | 0 | 1 | 1 | 1 | 1 | 0 | 1 | 1 | 0 | 1 | 1 | 1 | 1 | 1 |
| FHF<br>muted | 0 | 0 | 0 | 0 | 1 | 0 | 0 | 1 | 0 | 1 | 0 | 0 | 0 | 0 | 0 |
| FHF<br>state | 1 | 0 | 0 | 0 | 1 | 0 | 0 | 1 | 0 | 1 | 0 | 1 | 1 | 0 | 0 |

##### 4.2.1.2 Q-Learning based RL model

The Q-learning model is built with the state and action spaces defined by the cardiogenesis Boolean network model. For this model, FHF attractor state is considered to be the desired target state. The goal of the model is to find the optimal sequential gene expression modulations for reprogramming any of the other states of the Boolean network to FHF state.

The model training converged after about 25 episodes (see Figure 2). The coupled model learnt to transition from a starting state to the target state as the training progresses. This is evident from Figure 3, wherein at the initial episodes the model converges to the same intermediate attractor states multiple times consecutively (e.g. ∼23 times for Inactive attractor state: 000000000000000), while after about 25 episodes the consecutive occurrence of intermediate attractor states reduced to zero. Thus, implying that the model learnt the required modulation to transition out from the intermediate attractor state and towards the target attractor state.

**Figure 2:**
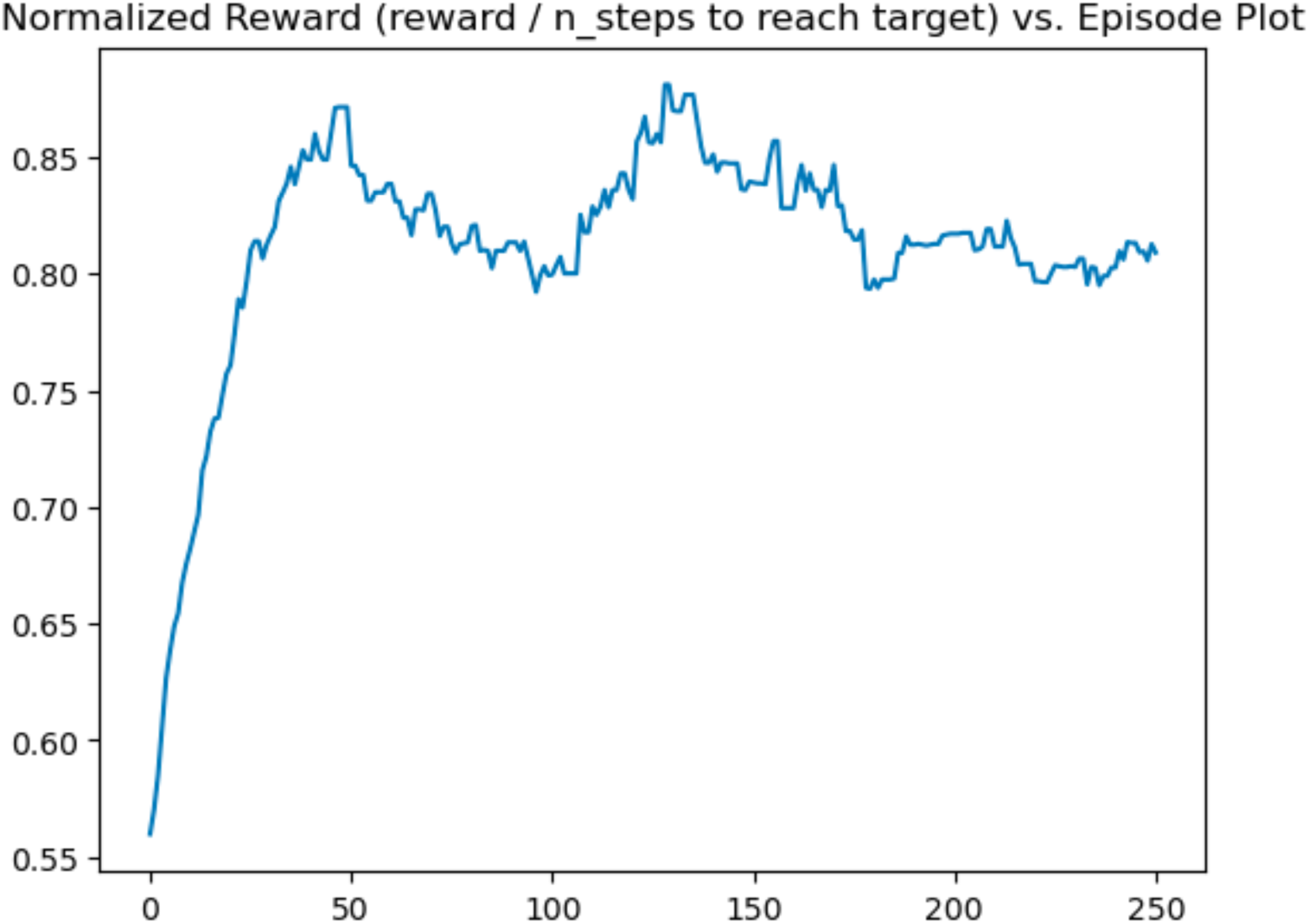
Convergence of the training of the cardiogenesis coupled model. Normalized cumulative reward (reward / no. of steps to reach target) in each episode is plotted against episode number

**Figure 3:**
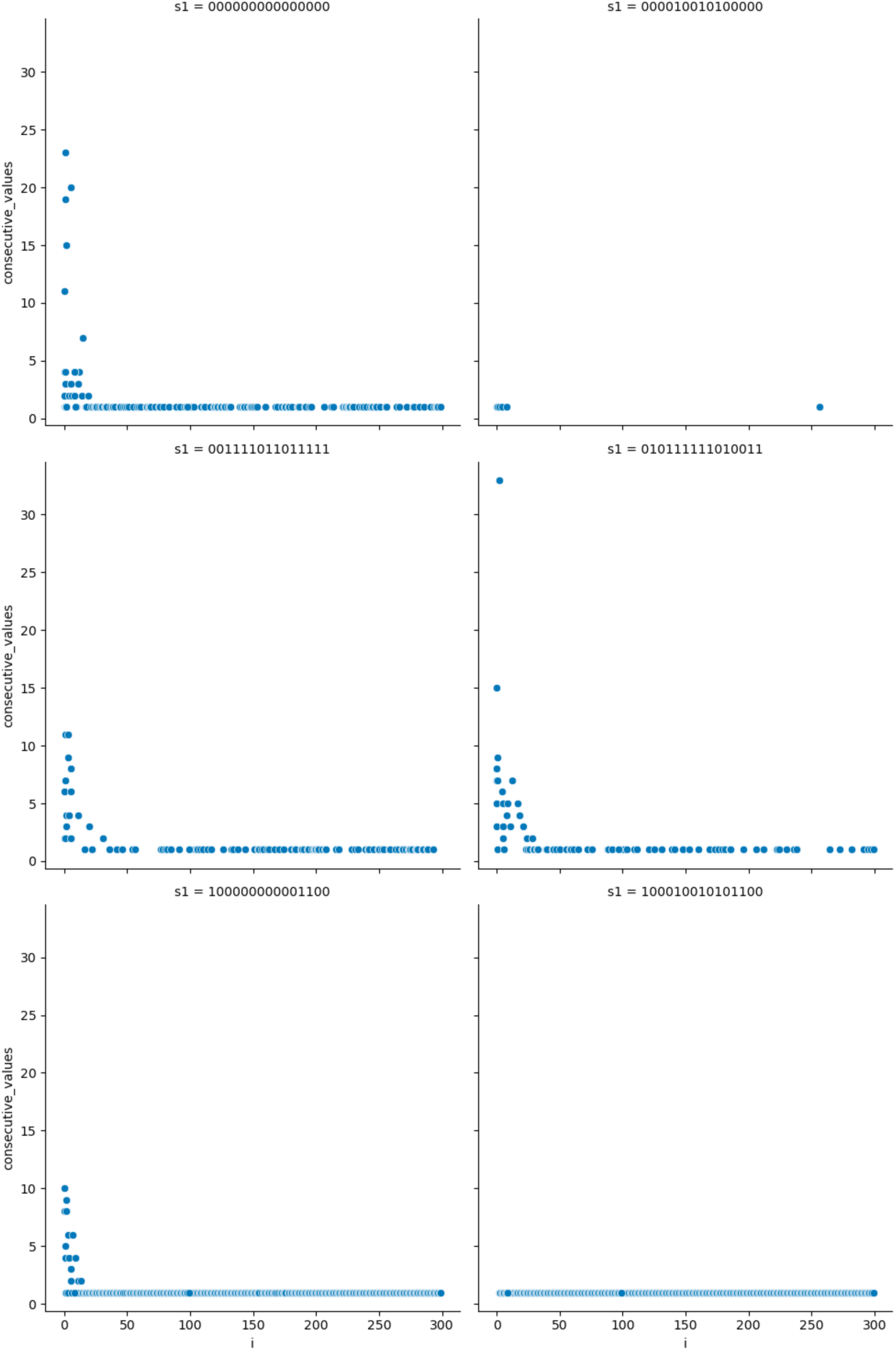
Number of times an attractor is visited consecutively as a function of episode numbers in cardiogenesis coupled model. s1: State vector of the attractors corresponding to each subplot; Inactive: 000000000000000; Active BMP: 100000000001100; SHF muted: 010111111010011; SHF state: 001111011011111; FHF muted: 000010010100000; FHF state: 100010010101100

#### 4.2.2 Coupled model predicts the required gene expression modulations for targeted reprogramming to FHF state

The learnt model (in the form of updated Q-table) was used to predict the set of gene expression modulations for converting any attractor state (except FHF state) to FHF state. The number of genes to be modulated and the details of their sequence are shown in Table 2. Out of 15 genes present in the network, only three genes are predicted to play a role in converting to FHF state. Moreover, the transition between various perturbed and attractor states during the conversion to the target state along with the sequence of gene expression modulations are shown in Figure 4. Transition from all the attractors except ‘FHF muted’ passes through the intermediate attractor, ‘Active BMP’, in agreement with the evidence that Bmp2 stays active in FHF state [60].

**Figure 4:**
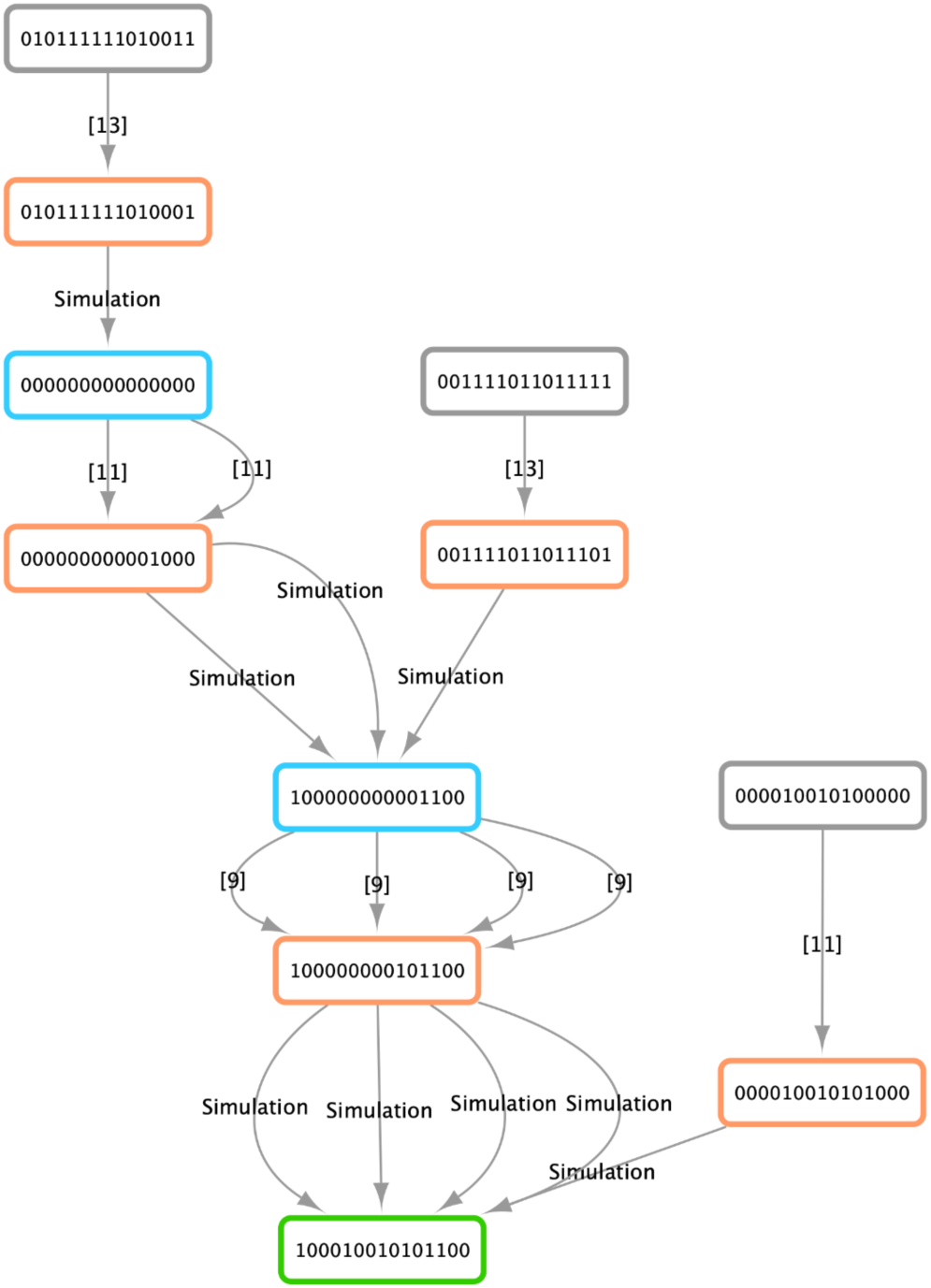
Required gene expression modulations and state transitions through starting attractors, perturbed states and intermediate attractors while converting an attractor state to FHF state in cardiogenesis coupled model. Nodes indicate Boolean network states and edges indicate their transition to other states either by gene expression modulation or by simulation of the state to its attractor; Node colors - Gray: Initial state, Orange: Perturbed state due to gene expression modulation, Blue: Initial or intermediate attractor states during transition to the target attractor state, Green: Target attractor state; Edge labels: Simulation: Simulate Boolean network model to find the attractor state corresponding to the current (perturbed) state, Values in square brackets indicate the modulated gene - [3]: Foxc1_2, [9]: Tbx5, [11]: exogen_BMP2_I, [13]: exogen_CanWnt_I; Attractor states mapping - Inactive: 000000000000000; Active BMP: 100000000001100; SHF muted: 010111111010011; SHF state: 001111011011111; FHF muted: 000010010100000; FHF state: 100010010101100

**Table 2:**
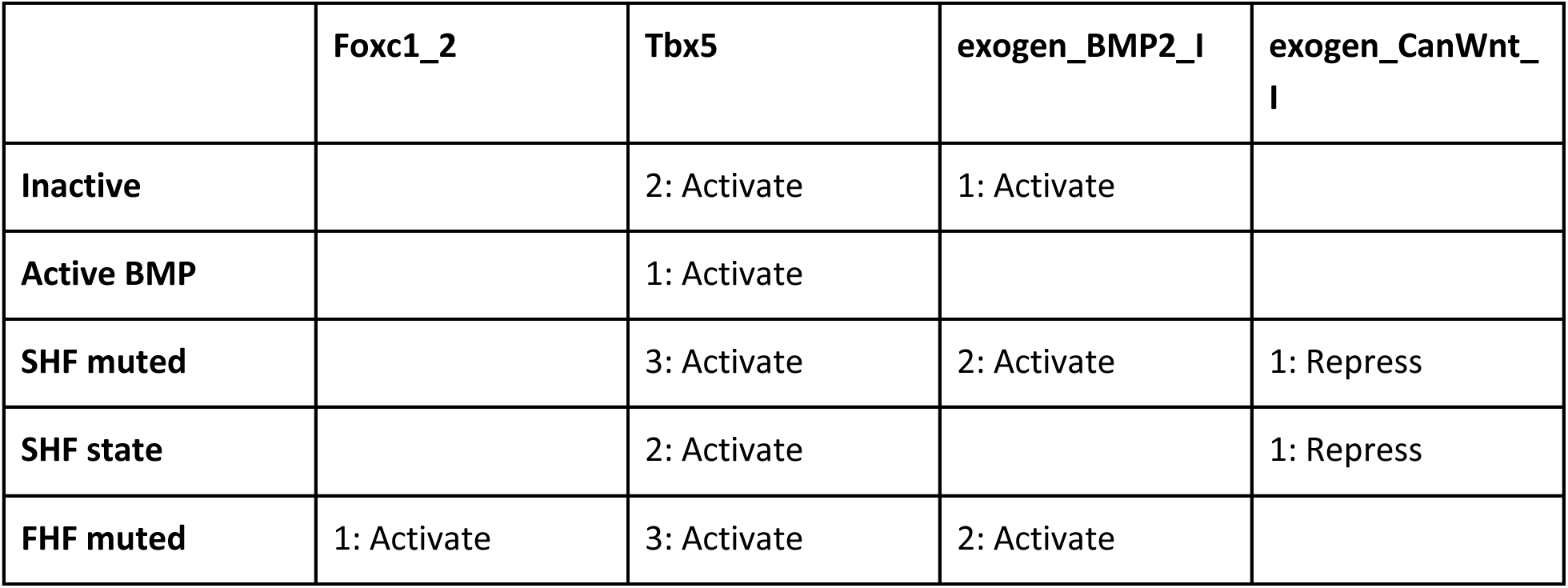
Gene expression modulations predicted by the proposed method for converting to ‘FHF state’ from any of the other attractor states in cardiogenesis coupled model. Rows: Starting attractor states; Columns: Genes to be modulated; Numeric values in the cells indicate the sequence of modulations

|  | Foxc1_2 | Tbx5 | exogen_BMP2_I | exogen_CanWnt_I |
| --- | --- | --- | --- | --- |
| Inactive |  | 2: Activate | 1: Activate |  |
| Active BMP |  | 1: Activate |  |  |
| SHF muted |  | 3: Activate | 2: Activate | 1: Repress |
| SHF state |  | 2: Activate |  | 1: Repress |
| FHF muted | 1: Activate | 3: Activate | 2: Activate |  |

The shortest sequence of gene expression modulation is one, and is required to convert ‘Active BMP’ attractor state to FHF state. The longest sequence is of three genes, and is required to convert ‘SHF muted’ state to FHF state. Moreover, the set of modulations required for converting SHF related attractors (i.e. SHF and ‘SHF muted’) to FHF states predicted the repression of exogen_CanWnt_I signal. This is in concordance with the evidence that exogen_CanWnt_I is required for SHF state [60], and thus, moving away from it required the repression of exogen_CanWnt_I.

### 4.3 Coupling RL with Boolean network model reduces the effective state space and makes the predictions biologically relevant

As detailed in section 3.2.2, Q-learning based RL model relies on and updates a state-action value table (Q-table) during model training to learn optimal actions for the states of the environment. With large state space, for example, 32,768 states for the cardiogenesis network, exploring all the states and all the actions in them can be combinatorially challenging (32,768 x 15 combinations in this case).

The Boolean network models allow predicting the attractor states that represent biological phenotypes. All the other (transient) states resulting from gene expression modulations or otherwise are known to transition to one of these attractor states eventually [31], [66]. Thus, in terms of the coupled model, any alteration in the state of the RL environment due to the action of RL agent, would eventually converge to an attractor state. This implies that the effective state space that needs to be considered by the RL model reduces from all the possible states to only the attractor states in a particular Boolean network model.

In the case of the cardiogenesis model, this amounts to a reduction of the number of states from 32,768 to 6 attractor states. Thus, instead of learning the gene expression modulations required for transitioning from each state in the state space to the target state, the model has to only learn to predict the modulations for transitioning a given attractor state to the target attractor.

Moreover, the ability to predict the effect of gene expression modulation directly on the dynamic behavior of the GRN ensures that the predictions made by the model are biologically relevant as long as the Boolean network model truly represents the underlying biological system dynamics.

## 5. Discussion

Ability to reprogram cells to desired cell types lie central to regenerative medicine. However, the complexity behind precise alteration of gene expression profiles of cells, and their subsequent reprogramming, demands approaches to *a priori* predict the required gene expression modulation. To this end, computational methods that can utilize available biological data and data-driven approaches concertedly hold the key. This study proposed one such computational method.

The proposed method captures the core dynamics of gene expression regulation with the help of a Boolean model of the concerned gene regulatory network, thereby, representing the core biological context. It further couples this model with Reinforcement Learning, a powerful approach for solving sequential decision-making tasks [67]; thus, allowing the proposed method to predict biologically relevant sequential gene expression modulations for targeted cellular reprogramming.

The relative simplicity of Boolean modeling formalism allows modeling biological networks with hundreds of components [31], and consequently, represent a larger context of biological dynamics. Moreover, recent development of computational methods for inferring Boolean network models from high-throughput data (e.g. single cell transcriptomic data) [68], [69], [70] makes it easy to develop such large-scale Boolean network models. Thus, the choice of Boolean modeling formalism in the proposed method makes it possible to scale the method to larger regulatory networks. Similarly, the choice of Q-learning algorithm for implementing the RL models also has several advantages: (1) being easily interpretable; (2) relatively less number of hyperparameters compared to other RL algorithms (e.g. Deep Q-learning), thus, easy to fine-tune; and (3) the simplicity of the algorithm makes training less time-intensive.

The proposed method relies on the Boolean network model for gaining the biological context. Thus, the method can be extended further towards: (1) incorporating Boolean models of genome-scale regulatory networks, thereby capturing system-wide biological context; and (2) using multi-omics data for developing such models, thereby capturing multilevel regulation of gene expression [35], [71], [72], as opposed to the current models that are derived only from transcriptomic data.

Furthermore, with the use of suitable rewards and penalties in the coupled models, the presented method offers a capability to incorporate additional biologically relevant constraints while predicting required gene expression modulations. For example, the method can be constrained to avoid transitioning to attractor states that are linked to undesirable phenotypes (such as tumor-promoting) while transitioning from starting state to the target attractor state.

## 5. Conclusion

To the best of the authors’ knowledge, this is the first attempt to integrate a Boolean network model with a RL model to solve a sequential gene expression modulations prediction problem, thereby, combining the strengths of available biological knowledge and a data-driven approach for gaining insight from it. The proposed method can benefit researchers in devising protocols for targeted cellular reprogramming that can overcome common pitfalls including low efficiency of conversion, and the risk of tumorigenicity and immunogenicity in the target cells. The method also opens ways to incorporate custom Boolean network models to capture the biological context of interest, and can be extended by incorporating other complementary omics datasets.

## CRediT authorship contribution statement

**Vivek Singh:** Conceptualization, Methodology, Software, Formal analysis, Investigation, Visualization, Writing - Original Draft, Writing - Review & Editing.

## Acknowledgements

The author would like to thank Thoughtworks for the support provided during this work. The author would also like to thank Avani Mahadik, Divye Singh, Janani Venugopalan, M Nimalan and Harshal Hayatnagarkar for interesting discussions and critical inputs on the study.

## Appendix: supplementary data

Following are the Supplementary data to this article.

