## Supplementary Data for "Optimizing sequential gene expression modulation for cellular reprogramming - Coupled Boolean modeling and Reinforcement Learning based method"

**Table S1: Nodes (genes) and their corresponding interactions represented as state transition functions for toy Boolean network model**

| Node | State transition function |
|---|---|
| x1 | x1 |
| x2 | x2 |
| x3 | x1 & !x2 |
| x4 | x3 \| x4 |

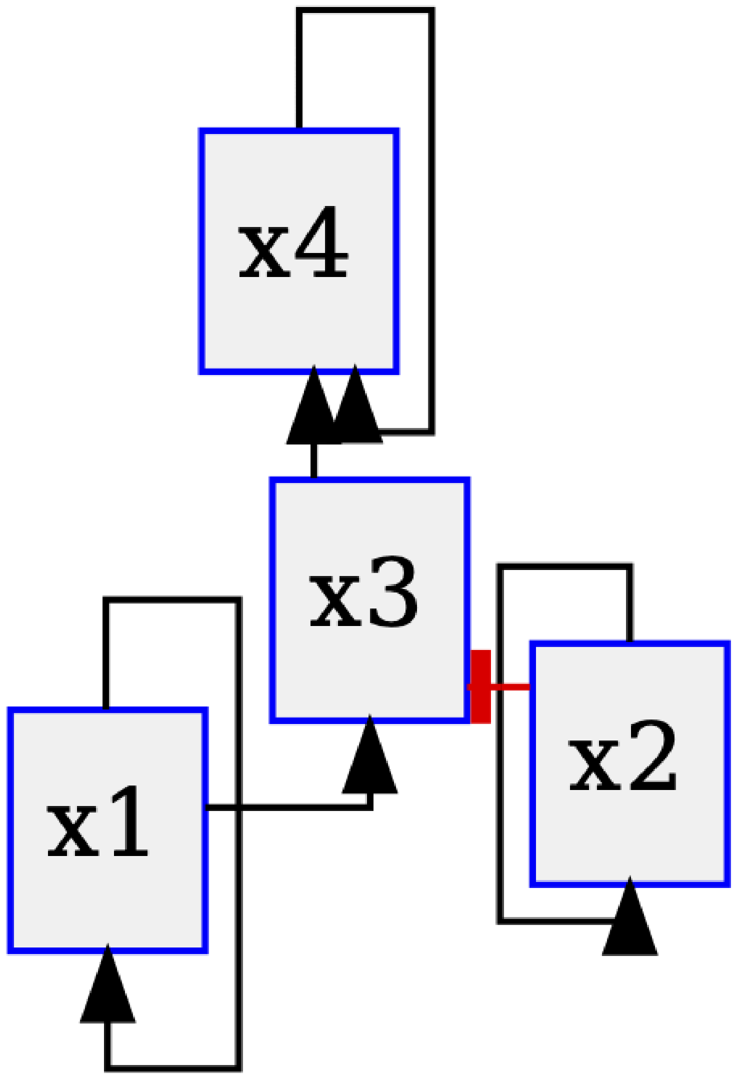


**Figure S1: Interaction graph of toy Boolean network model**

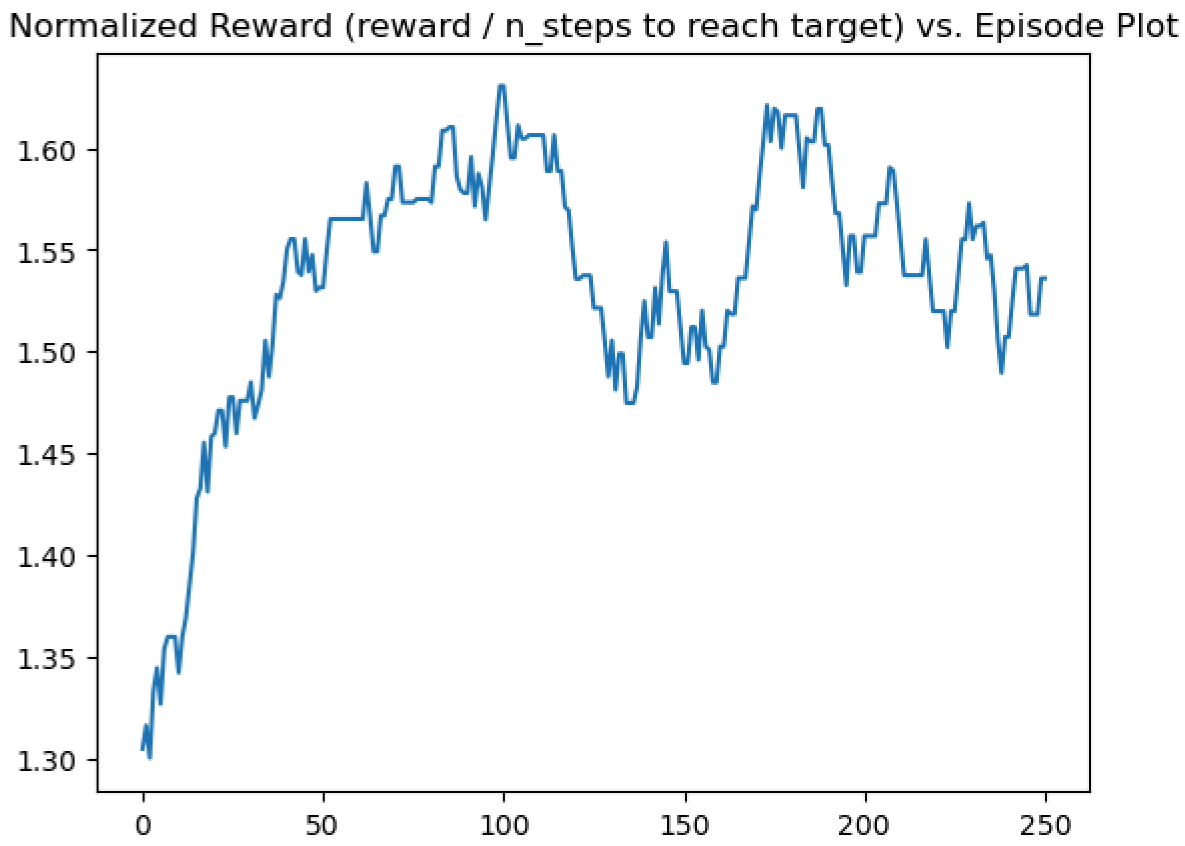


**Figure S2: Convergence of the training of the toy coupled model.** Normalized cumulative reward (reward / no. of steps to reach target) in each episode is plotted against episode number

**Table S2: Gene expression modulations predicted by the proposed method for converting to the attractor state '1011' from any of the other attractor states in the toy coupled model**

Rows: Starting attractor states; Columns: Genes to be modulated; Numeric values in the cells indicate the sequence of modulations

| | **x1** | **x2** |
|---|---|---|
| **0000** | 1: Activate | |
| **0001** | 1: Activate | |
| **0100** | 1: Activate | 2: Activate |
| **0101** | 2: Activate | 1: Activate |
| **1100** | | 1: Repress |
| **1101** | | 1: Repress |

**Table S3: Nodes (genes) and their corresponding interactions represented as state transition functions for cardiogenesis Boolean network model**

| Node | State transition function |
|---|---|
| exogen_BMP2_I | exogen_BMP2_I |
| exogen_BMP2_II | exogen_BMP2_I |
| exogen_CanWnt_I | exogen_CanWnt_I |
| exogen_canWnt_II | exogen_CanWnt_I |
| canWnt | exogen_canWnt_II |
| Bmp2 | (exogen_BMP2_II & !canWnt) |
| Foxc1_2 | (canWnt & exogen_canWnt_II) |
| Mesp1 | (canWnt & !exogen_BMP2_II) |
| Dkk1 | ((canWnt & !exogen_BMP2_II) \| Mesp1) |
| Tbx1 | Foxc1_2 |
| Fgf8 | ((Foxc1_2 & !Mesp1) \| (Tbx1 & !Mesp1)) |
| Isl1 | (((Tbx1 \| Fgf8) \| (canWnt & exogen_canWnt_II)) \| Mesp1) |
| GATAs | ((Tbx5 \| Mesp1) \| Nkx2_5) |
| Nkx2_5 | (((((Mesp1 & Dkk1) \| Tbx1) \| (Bmp2 & GATAs)) \| (Isl1 & GATAs)) \| Tbx5) |
| Tbx5 | (((Nkx2_5 & !((Tbx1 \| (Dkk1 & (!Tbx5 & !Mesp1))) \| canWnt)) \| (Mesp1 & !((Tbx1 \| (Dkk1 & (!Tbx5 & !Mesp1))) \| canWnt))) \| (Tbx5 & !((Tbx1 \| (Dkk1 & (!Tbx5 & !Mesp1))) \| canWnt))) |

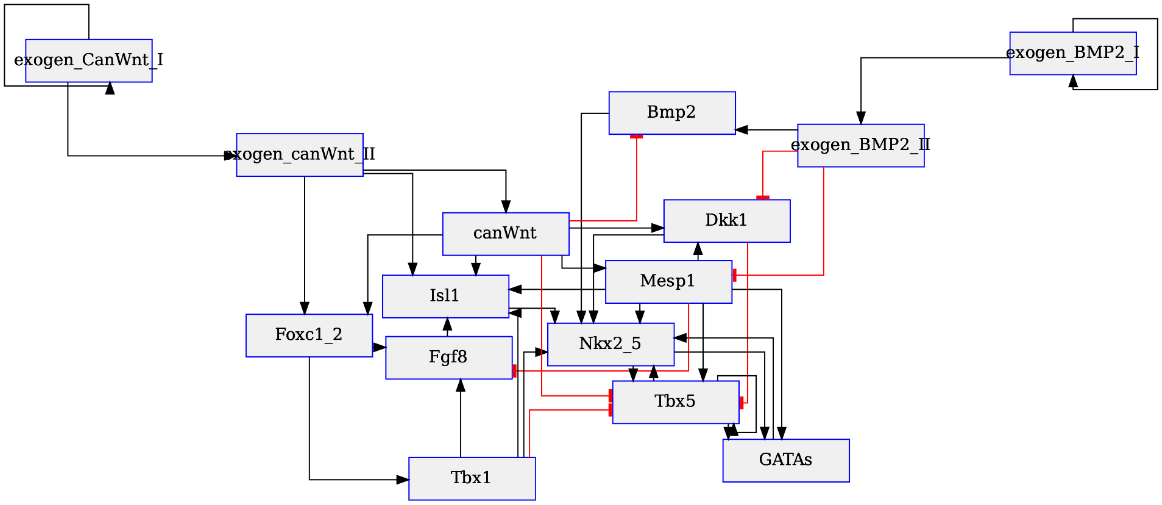


**Figure S3: Interaction graph of cardiogenesis Boolean network model**
